# Dynamics of hinged wings in strong upward gusts

**DOI:** 10.1101/2022.08.05.502962

**Authors:** Jonathan P. J. Stevenson, Jorn A. Cheney, James R. Usherwood, Richard J. Bomphrey, Shane P. Windsor

## Abstract

A bird’s wings are articulated to its body via highly mobile shoulder joints. The wings possess an impressive range of motion, which allows for movements that can modulate the production of aerodynamic load in flight. This is crucial for stability and control—particularly in gusty or turbulent environments. In this study, we develop a dynamics model to examine how a bird-scale gliding aircraft can use wing-root hinges (analogous to avian shoulder joints) to reject the initial impact of a strong upward gust. Our idea requires that the spanwise centre of pressure and the centre of percussion of the hinging wing start, and stay, in good initial alignment (the centre of percussion here is related to the idea of a ‘sweet spot’ on a bat, as in cricket or baseball). We propose a method for achieving this rejection passively, for which the essential ingredients are: (i) well-considered lift and mass distributions; (ii) constant-torque hinges; and (iii) a wing whose sections stall softly. When configured correctly, the gusted wings will first pivot on their hinges without disturbing the fuselage of the aircraft, affording time for other corrective actions to engage. We expect this system to enhance the control of aircraft that fly in gusty conditions.

## 1 Introduction

Flight in the low atmosphere (< 1000 m) is challenging and often dangerous. Aircraft and animals must contend with strong winds and gusty flows (Etkin 1981; Watkins *et al.* 2006) while steering clear of obstacles, terrain and other airborne objects. Birds are light and fly relatively slowly; they are especially vulnerable as windspeeds rise (Tennekes 2009). Yet many of them, noticeably the gilders, manage to fly with impressive agility and control in this realm of the atmosphere, their wings and tails tilting and flexing in response to the unsteady wind (Reynolds *et al.* 2014). Aircraft designers are now seeking to harness the principles of avian gust rejection for use in flight control systems (Ol *et al.* 2008; Wilson *et al.* 2019).

In this study, we focus on a specific idea: the utility of wing-root hinges for gust rejection on aircraft, particularly small unmanned aerial vehicles (UAVs). The work was motivated by a laboratory experiment in which a gliding barn owl responded to strong upward gusts (with a magnitude of 40-70% of the flight speed, and a width greater than the wingspan) by immediate rotation of its wings around the shoulder joints (Cheney *et al.* 2020). The motion of the wing masses effectively absorbed the sudden impulse from the extra aerodynamic load, allowing the torso and head, together equivalent to a fuselage, to maintain a smooth flight trajectory. Cheney *et al.* named this effect *inertial rejection;* it relies purely on the displacement of wing inertia and does not require active control.

Inertial rejection can be enhanced by exploiting an intrinsic property of hinged wings and, for that matter, all rigid (or near-rigid) pivoting masses: the *centre of percussion.* It is that point on the mass at which a sudden transverse load does not transmit any immediate reaction force through the pivot. In other words, the pivot will not ‘feel’ anything in the transverse direction. Players of bat-and-ball sports, particularly cricket and baseball, will have practical familiarity with the centre of percussion, as it is closely related to the so-called *sweet spot*—that special zone on a bat where a ball can be struck without causing unpleasant jarring of the hands (Brearley *et al.* 1990; Cross 1998). In general, the centre of percussion *P* of a hinged object, of mass *m,* is located at a distance (supplemental information (SI) section 6)

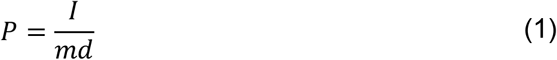

from the pivot point. Distance *d* is that from the hinge to the centre of mass of the object, and *I* the corresponding mass moment of inertia about the hinge.

As such, hinged wings can perform a similar role to suspension systems on terrestrial vehicles. Sabins (1937) patented an early imagining of suspension for light aircraft, in which each (rigid) wing was hinged to the fuselage on a root pin joint and supported on a shock-absorbing strut. The wings were to deflect up or down in response to changing air loads in flight, and to shocks during landing, with the aim of reducing structural stresses and enhancing passenger comfort. More recently, studies have suggested that wing-root hinges can improve the gust response of small-scale aircraft (Webb & Costello 2008; Stewart *et al.* 2008) and may even provide opportunities for tuning flight stability and performance (Paranjape *et al.* 2012; Leylek & Costello 2015). To our knowledge, though, gust-rejecting hinged wings have not yet seen mainstream application on commercial aircraft at any scale, nor has the centre of percussion been used to enhance any of the designs that do exist.

In this study, we develop a dynamics model to examine how the action of a wing-root hinge can mitigate the initial impact of an upward gust (hereafter *upgust*) on the fuselage of a small gliding aircraft. The gust is wider than the wingspan of the flyer, as in the owl experiment of Cheney *et al.* (2020). Having explored the nature of the centre of percussion and its basic role in the transmission of load from the hinging wings to the fuselage, we propose a method to exploit the mechanics for immediate passive gust rejection. Altogether, the results inform the design of novel wing suspension systems for aircraft that require smoother flight, *e.g.,* those carrying fragile payloads, cameras or sensors in difficult conditions.

## 2 Methods

The modelled system is a mechanical analogue for a bird, consisting of a central fuselage (torso) mass and two rigid wing beams (Figure 1). Each wing root is hinged to the fuselage on a pin joint. The fuselage is free to translate along a vertical line, while the wings can rotate on their hinges in symmetry, *i.e.,* wing motion is mirrored about the vertical centreline (this ‘reduced order’ representation is based on the initial kinematic response of the owl to an upgust, in which the wings rotated symmetrically about the shoulders without significant morphing). As such, the system has two degrees of freedom; we choose fuselage height *z* and wing angle *θ* (from the horizontal) as the most convenient generalised coordinates to describe the dynamics.

**Figure 1.**
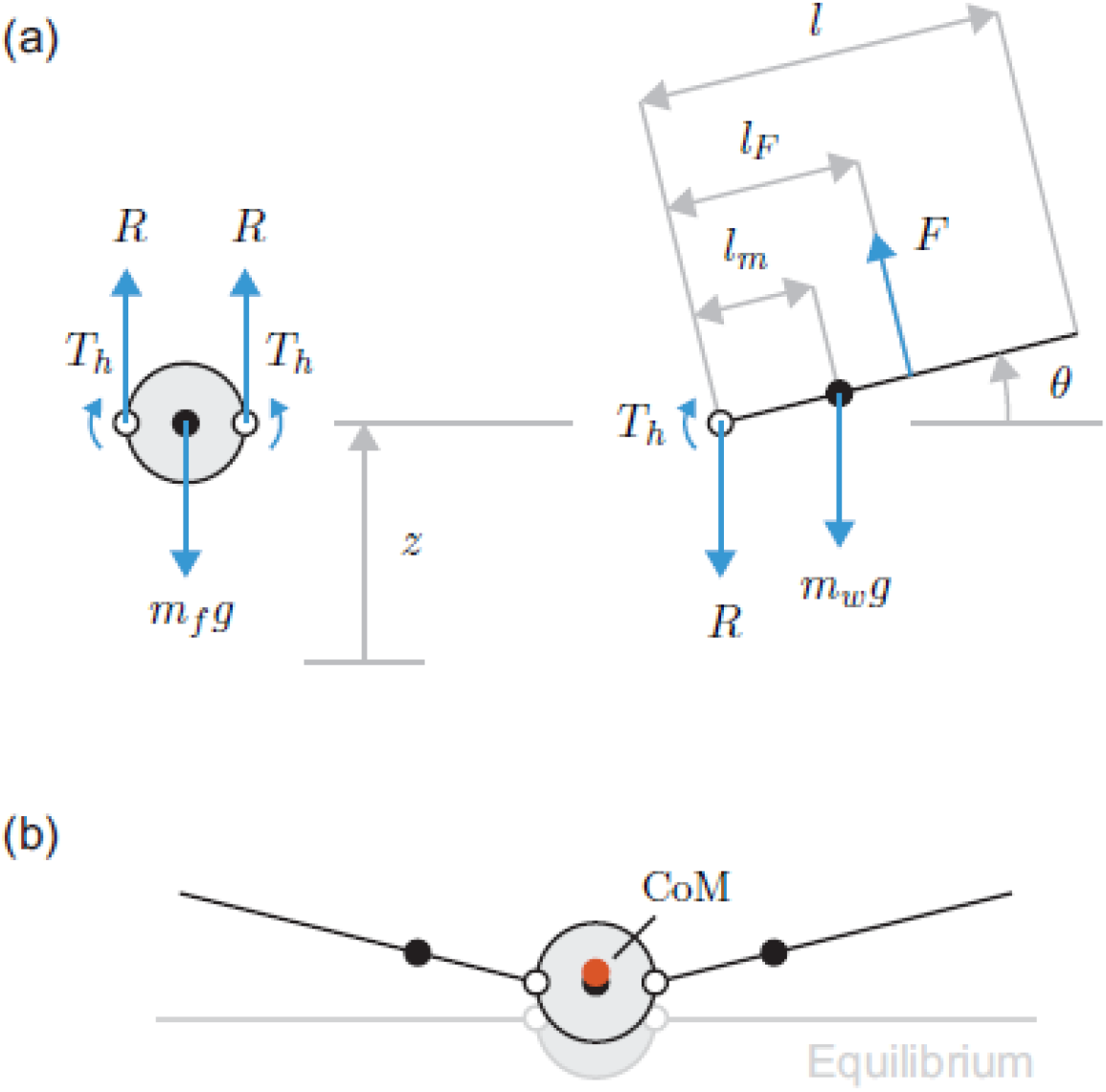
The modelled system. (a) Free-body diagrams for the fuselage (left) and wing (right) masses, horizontal forces omitted (they do not affect the dynamics). The fuselage can translate vertically, while each wing can hinge about its root end on a pint joint (open circle). The fuselage is regarded as dimensionless, and the aerodynamic force upon it is neglected. (b) Motion occurs when extra vertical force on the wings pushes the system away from equilibrium. The centre of mass (CoM) of the overall system (orange circle) lies, by definition, between those of the wings and that of the fuselage (all black circles).

At first, the system travels in level equilibrium (lift equals system weight) with constant forward flight speed *U.* The initial relative wind is therefore horizontal, also with magnitude *U.* The system then encounters a strong upgust (whose width far exceeds the wingspan) that produces extra lift on the wings, impelling them upwards, and the fuselage translates as load is transmitted to it via the hinges. In this study, we seek to understand these vertical motions, particularly at the level of the fuselage.

We simplify the problem by neglecting drag and pitch. For the changes in angles of attack we consider, the upward component of wing drag is small enough not to affect the vertical motion of the system, and the pitching moment is assumed not to change throughout. The latter assumption requires that the wing be hinged at its aerodynamic centre (the chordwise point about which the pitching moment is practically constant with incidence). Further, the system has no tail, so there is no other source of pitching moments.

Figure 1 shows free-body diagrams for the fuselage and a single wing, horizontal forces omitted. The relevant equations of motion for this system are (see SI section 1)

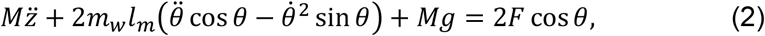

which derives from a balance of forces on the overall centre of mass (CoM), and

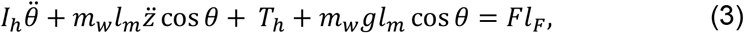

which is a moment sum on each wing about its hinge. See Table 1 for symbol definitions. Note that distance *l_m_* is a function of the wing mass distribution, as is the mass moment of inertia *I_h_.* On the right-hand side of the equations, *F* is the component of wing lift in the plane of motion (the page), which acts at distance *l_F_* from the hinge.

**Table 1.** Properties and inputs of the modelled system. †Stiffness is varied later.

| Symbol | Description | Value | Unit |
| --- | --- | --- | --- |
| $g$ | Magnitude of gravitational acceleration | 9.81 | m/s<sup>2</sup> |
| $m_f$ | Mass of fuselage | 0.25 | kg |
| $m_w$ | Mass of single wing | 0.025 | kg |
| $M$ | Mass of overall system ($= m_f + 2m_w$) | 0.3 | kg |
| $\mu_f$ | Mass fraction of the fuselage ($= m_f/M$) | 0.8<sup>†</sup> | - |
| $\mu_w$ | Mass fraction of the wings ($= 2m_w/M$) | 0.2<sup>†</sup> | - |
| $c$ | Chord length of wing section | 0.15 | m |
| $l$ | Spanwise length of wing | 0.4 | m |
| $l_m$ | Centre of mass of wing from hinge | 0.133 | m |
| $I_h$ | Mass moment of inertia of wing about hinge | 6.67e-4 | kg m<sup>2</sup> |
| $P$ | Centre of percussion of wing from hinge | 0.2 | m |
| $k_t$ | Torsional hinge stiffness | 0<sup>†</sup> | N m/rad |
| $\rho$ | Air density | 1.2 | kg/m<sup>3</sup> |
| $U$ | Forward flight speed ($=$ relative wind) | 8.0 | m/s |

We model the ‘total’ torque at each hinge as having two parts, or *T_h_* = *T*_*h*0_ + Δ*T_h_.* The static part *T*_*h*0_ is that necessary to keep the wings in level equilibrium for ordinary flight. It is found from the combination of equations (2) and (3) at equilibrium:

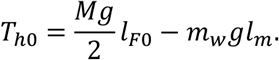

For this system, the static torque for level equilibrium is restorative in sense, *i.e.*, pulling each wing downwardly (*l_m_* < *l*_*F*0_ and *m_w_* ≪ *M* from Table 1). This is also true for birds (Gray 1968). The dynamic part Δ*T_h_* only engages when the wings deviate from level equilibrium. It depends on the mechanics of the hinge—whether they are passive, active or some combination of both. In section 3.5, we consider the effect of a linear torsional spring with restorative behaviour (Δ*T_h_* = *k_t_θ*, where *k_t_* is the stiffness). Such a hinge is totally passive, and can also provide the necessary static torque by pre-torsion to the correct angle.

Reaction *R,* present as an equal-and-opposite force pair at each hinge (Figure 1), determines the motion of the fuselage. The equation of motion is

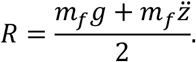

Note that *R* has static (*R*_0_ = *m_f_g*/2) and dynamic 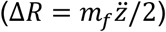 parts. The dynamic part Δ*R* governs the response of the fuselage to the gust.

Frequent comparison is made in later sections to a system with fixed wings, *i.e.,* a rigid aircraft without wing hinges. Its equation of motion is

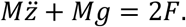

In this case, the wings and fuselage must move in tandem. The point of application of force *F* therefore does not matter.

### 2.1 Linearisation

The equations of motion for the hinged system are nonlinear, second-order and coupled by the generalised coordinates. We now linearise them about *θ* = 0 to simplify the mechanics of the problem for clearer insight on the basic mechanics. This does limit the results to |*θ*| ≲ 20 degrees (when the small-angle approximation cos*θ* ≃ 1 passes 5% error), but the percussion effect is expected to be most applicable at low angles anyway.

Using the small-angle approximation (cos*θ*~1) and the fact that the initial angular wing velocity is small, especially when squared 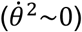, we find the linearised equations of motion,

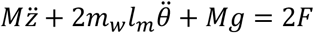

and

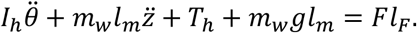

One further simplification—that total load be regarded as the sum of static (subscript 0) and dynamic (Δ) parts—allows equilibrium loads to be subtracted from these equations. Therefore (see SI section 2),

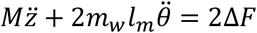

and

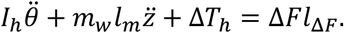

Force *F* (hence Δ*F*) now effectively points upwards at all times.

### 2.2 System parameters

The size and overall mass of the system are based on the owl in Cheney *et al.* (2020). Some parameters have been rounded for convenience (Table 1). As such, we are strongly inspired by the dynamics of the owl, but do not intend to mimic all its aeromechanical complexities.

For the spanwise wing mass distribution, we choose a linear (triangular) function. This is a convenient approximation for the wings of birds (see van den Berg & Rayner (1995), for example) and aircraft alike. For a wing of length *l,* it has the form (see SI section 3)

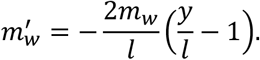

*y* is the spanwise coordinate, running from root to tip. The centre of percussion of this distribution lies at *l*/2, which, as discussed shortly, aligns perfectly with the centre of pressure for our chosen lift distribution at equilibrium. Cheney *et al.* (2020) found that such equilibrium alignment is *almost* true for the barn owl—within just 0.067*l*. Here, we examine this case of perfect alignment because it may provide the largest potential gust rejection benefit.

### 2.3 Aerodynamics formulation

As described, *F* is the aerodynamic force on each wing in the (vertical) plane of motion. Our aerodynamics formulation models this force using a blade-element approach.

The upward projection of *F* (*i.e.*, *F* cos*θ*) determines the vertical motion of the system, and, in the linearisation of *θ,* is approximately equivalent to *F* itself. All subsequent references to *F* therefore refer to this vertical projection. As soon discussed, the tested gust also allows for linearisation of the dynamic angle of attack Δ*α*. These linearisations together simplify the formulation and facilitate basic insight. There are several assumptions:

i. The wing planform is rectangular and untwisted. (That it has a triangular mass distribution is explained by a variation in material density.)
ii. The flow is two-dimensional. (Note that the effect of spanwise flow reduces with *θ*.)
iii. All wing sections are symmetrical about the chord line and have identical aerodynamic properties.
iv. Aeroelastic flexure (of the wing structure) is absent.
v. The effect of apparent air mass is negligible.
vi. Aerodynamic force acts in a quasi-steady manner.

With respect to point (vi), we recognise that aerodynamic force actually takes time to ‘build up’ in response to gusts and other wing motions (Fung 1993; Wright & Cooper 2007). We checked this effect using classical approaches, viz. Küssner and Wagner lift functions (SI section 5), and found that it does indeed delay the gust response of the system, effectively pushing the quasi-steady results back in time (the shape and character of the response is otherwise very similar). We therefore decided to neglect ‘build-up’ in the model proper and to employ the quasi-steady formulation throughout. The main formulae of this formulation now follow (see SI section 4 for details).

Force *F* is the integral resultant of the spanwise aerodynamic lift distribution, denoted *F*′(*y*). Linearised, this distribution is

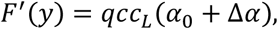

where *q* = *ρU*^2^/2 is the dynamic pressure, and *c* the (constant) chord length. Recall that all wing sections are identical and have the same lift curve, so the lift coefficient distribution *c_L_* depends only upon: (i) the static angle of attack *α*_0_, necessary for weight support at equilibrium; and (ii) the dynamic angle of attack increment Δ*α*(*y*), which embodies any changes brought about by the gust and/or motion of the system. We find that (SI section 4)

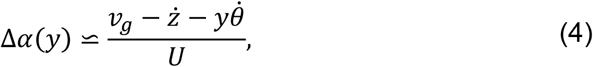

where *v_g_* is the gust velocity (introduced shortly). The total force on each wing is then

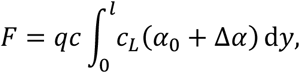

and its moment about the hinge axis

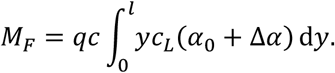

Quotient *M_f_/F* = *l_F_* then defines the spanwise point of action or *centre of pressure* of the force. In our calculations, we use discretised versions of these integrals with 50 spanwise points along each wing.

Velocity *v_g_* describes the upward gust intensity in the direction of flight. We use the standard ‘1 – cosine’ profile from the FAA airworthiness regulations (FAA 2020),

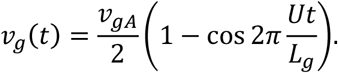

Here, *v_gA_* is the peak velocity, and *L_g_* the physical length, of the gust (Table 2). The gust is otherwise homogenous in the spanwise direction, *i.e.,* an unlimited spanwise ‘extrusion’ of this profile. In this study, we test a nominal peak gust velocity of 30% forward speed, or 0.3*U* (by nominal, we mean that the Küssner function has not been applied). Such a gust would seriously challenge the flight of most animals and small aircraft, but is still modest enough not to undermine the assumption of linearity (in *θ*) for the initial response. Indeed, upward acceleration of the wings is restrained by their mass.

**Table 2.** Properties of the vertical gust, also based on Cheney *et al.* (2020).

| Symbol | Description | Value | Unit |
| --- | --- | --- | --- |
| $v_{gA}$ | Peak gust velocity | 2.4 | [m s<sup>-1</sup>] |
| $L_g$ | Gust length along flight direction | 1.4 | [m] |

At equilibrium, there is no gust or wing motion, so Δ*α* = 0 everywhere. The force distribution *F*‱ is therefore rectangular, and its resultant *F*_0_ lies halfway along the wing (*l*/2). This is clearly an approximation, but it is an effective way, along with a triangular mass distribution, to put the centres of pressure and percussion into perfect alignment at equilibrium.

Temporal solution of the equations of motion was carried out in MATLAB® (MathWorks, Massachusetts) using the ODE45 Runge-Kutta solver. At each time point, the aerodynamic loads were first computed from the current state variables before being passed to the equations for solution of the next iteration. The algorithm automatically adjusted the size of the time step according to local gradients in the solution, but for added robustness we specified a maximum step of one-sixth of the gust duration (*L_g_*/6*U* or ~30 ms). Once finished, ODE45 returned a solution sampled at the requested time points—in this case, every 5 ms.

## 3 Results

First, we develop the basic mechanics of the centre of percussion for point force (section 3.1). We then extend the analysis to distributed aerodynamic loading, considering two lift curves: one with constant slope (section 3.2) and another with ‘soft stall’ behaviour (section 3.3). We also explore the effect of relatively heavy wings (section 3.4). Finally, we introduce dynamic hinge torque (section 3.5) via linear torsional spring stiffness.

### 3.1 Basic mechanics

Hinges allow the wings to rotate in response to external impulses, including gusts. Consider first a system with hinges that each produce a constant torque, equal only to the static value (*T_h_* = *T*_*h*0_). The system therefore supports level flight *exactly.* If either wing is struck by a transverse perturbing force Δ*F,* an associated reaction increment Δ*R* may develop, or be *transmitted,* to the fuselage via the corresponding hinge (we omit *increment* from now on). The underlying mechanics are of course embodied within the equations of motion, and from the linearised forms we can derive a concise expression that connects Δ*F* to Δ*R.* It is given by (see SI section 6)

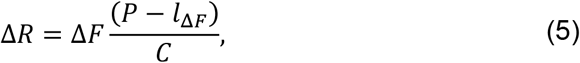

where *l*_Δ*F*_ is the distance from the hinge to the force Δ*F, P* is the position of the centre of percussion on the wing, and *C* = *P* – (*μ_w_/μf*)(*l_m_* – *P*) is a positive constant. The formula works best at moderate wing angles (|*θ*| ≲ 20 degrees, at which the approximation cos*θ* ≃1 crosses 5% error) and applies during the crucial early moments of the response.

According to the expression, it is the position and magnitude of force Δ*F* that govern the initial transmission of reaction Δ*R* to the fuselage. Static loads do not matter. Assuming, for now, a constant point force, there are three possible scenarios: (i) inboard or ‘armpit’ loading, in which Δ*F* acts inside *P.* Immediate upward reaction develops on the fuselage, with a magnitude that depends on the degree of misalignment with *P*; (ii) outboard loading, in which Δ*F* outside *P.* Reaction now develops in the downward direction, whose magnitude also depends on the misalignment; or (iii) aligned loading, in which Δ*F* acts *at P.* The bracketed term vanishes and no immediate reaction develops. We call this the *percussion effect,* and it works regardless of the magnitude of ΔF. Note also that, if the reaction is nullified, so too is the rolling moment it would otherwise apply to the fuselage (particularly important when the wings are loaded asymmetrically and these moments do not balance out).

For the case of fixed wings, Δ*R* = *μ_f_*Δ*F* (see SI section 6). Analytical comparison between the fixed and hinged cases (SI section 7) reveals that the fuselage experiences less absolute reaction in the latter, provided Δ*F* acts within a specific interval on the wing: *P* ±(*P* − *μ_w_l_m_*). This interval is symmetrical about *P* and its width depends on the wing mass distribution. For a triangular distribution, it is quite wide, covering the central ~89% of the wing length. The simple process of hinging therefore modulates the transmission of reaction to the fuselage for a wide range of loading points, while optimal tuning (Δ*F* at *P*) eliminates it altogether. Figure 2 provides a complete summary.

**Figure 2.**
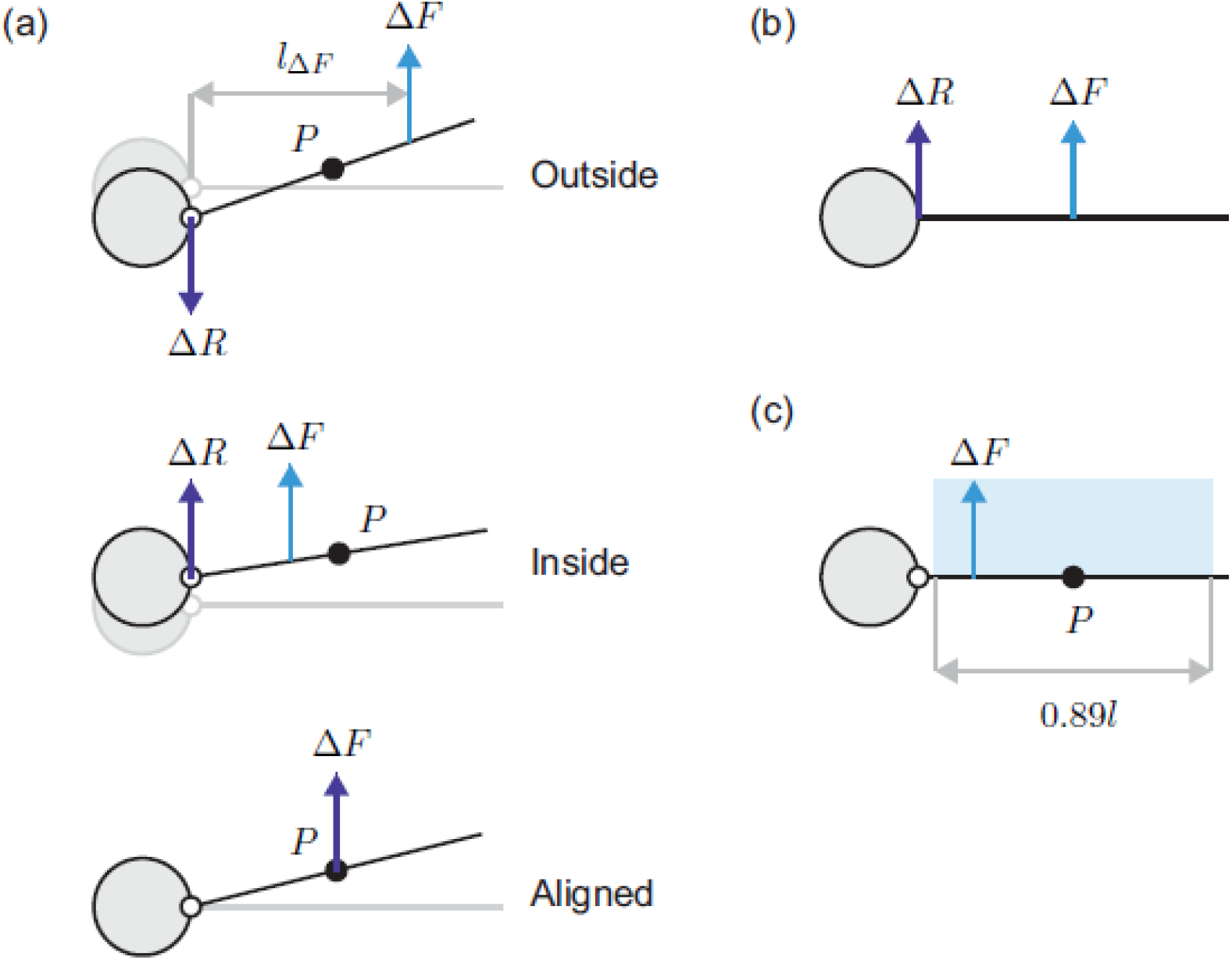
‘Pseudo’ free-body diagrams of load transmission from wing to fuselage. Hinge torque is constant. (a) Linearised dynamics of the hinged wing. The direction of the fuselage reaction increment Δ*R* varies with the spanwise location of applied force ΔF. Force through *P* produces no immediate reaction. (b) Dynamics of the fixed wing. Δ*R* acts in the same direction as Δ*F* with magnitude Δ*R* = *μ_f_*Δ*F*. Note that Δ*R* < Δ*F* because *μ_f_* < 1, *i.e.,* some of the applied force is always required to accelerate the wing mass. (c) A hinged wing with triangular mass transmits less reaction to the fuselage than a fixed one, provided Δ*F* acts within the wide, symmetrical interval (blue, to scale) centred on *P.*

Note that force Δ*F* (2Δ*F* with two wings) also applies an upward impulse to the system *as a whole,* and unless this is somehow countered by opposing action, the system will drift away from equilibrium indefinitely. Fortunately, aerodynamic damping prevents this in practice, but alone this is a relatively slow process. Proper control authority demands something much faster. This is an important point, revisited later when we discuss *aerodynamic rejection.*

### 3.2 Upgust response

The basic mechanics show that unless transverse perturbing force acts at the centre of percussion of the wing, sudden vertical reaction will be transmitted to the fuselage. Now, in real gusts, the perturbing force does not act solely at one point; rather, it is the integral resultant of any extra, distributed aerodynamic load across each wing. In order to realise the percussion effect in this case, we could align the centres of pressure and percussion at equilibrium, as we have already, and then strive to *keep them together* as the wing rotates under the action of the gust. We soon consider a method for doing this; first, we develop some analytical prerequisites. Hinge torque will remain constant (*T_h_* = *T*_*h*0_) until specified otherwise.

At equilibrium, the force distribution across each wing is constant. All wing sections operate at the same point on the lift curve. Consider first that each wing section has the same, symmetrical *linear* lift curve (LLC), or

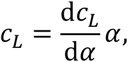

where d*c_L_*/d*α* is the (constant) slope. As such, lift varies in proportion to the angle of attack, and there is no stall in either direction. This is the usual choice when modelling minor gusts (Wright & Cooper 2007), and for the description of the extended linear region in real, dynamic stall (McCroskey 1981). When first encountered by the aircraft, the upgust begins to incline the relative wind vector from its equilibrium orientation, pushing the angle of attack of all wing sections up the lift curve. This occurs uniformly across the wing at first, scaling the equilibrium lift distribution such that its centroid, or centre of pressure, is momentarily preserved. As gust force builds, the wing is duly impelled to rotate upwards. This induces a distribution of relative downward flow *v_y_* (hereafter *relative down-flow*) across the wing—strongest at the tip, where the linear wing speed is highest. The relative downflow opposes the gust velocity *v_g_* and acts to temper the rising angle of attack Δ*α,* according to equation (4):

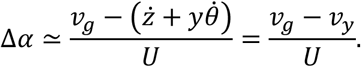

Here, *v_y_* is the local relative down-flow at distance *y* from the hinge, equal and opposite to the absolute vertical speed of the wing section at that location 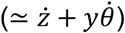. The presence of this relative down-flow gradient means that the spanwise force distribution along each wing is now skewed. The centre of pressure *l_F_* therefore drifts inboard from its equilibrium position and away from the centre of percussion. The force *increment* drifts, too, and equation (5) promises that reaction on the fuselage will inevitably result.

Figure 3 shows the impact of the tested gust (Table 2) on three systems: (i) entirely immobile, in which the wings and fuselage are both clamped in their equilibrium positions throughout; (ii) fixed-wing; and (iii) hinged-wing. Comparison between (i) and (ii) reveals the tempering effect of relative down-flow on the applied force (Figure 3a). The hinged wing experiences this to an even larger extent, being lightweight and better able to retreat from the gust than the fixed system (whose *full* inertia must be overcome for wing motion to begin).

**Figure 3.**
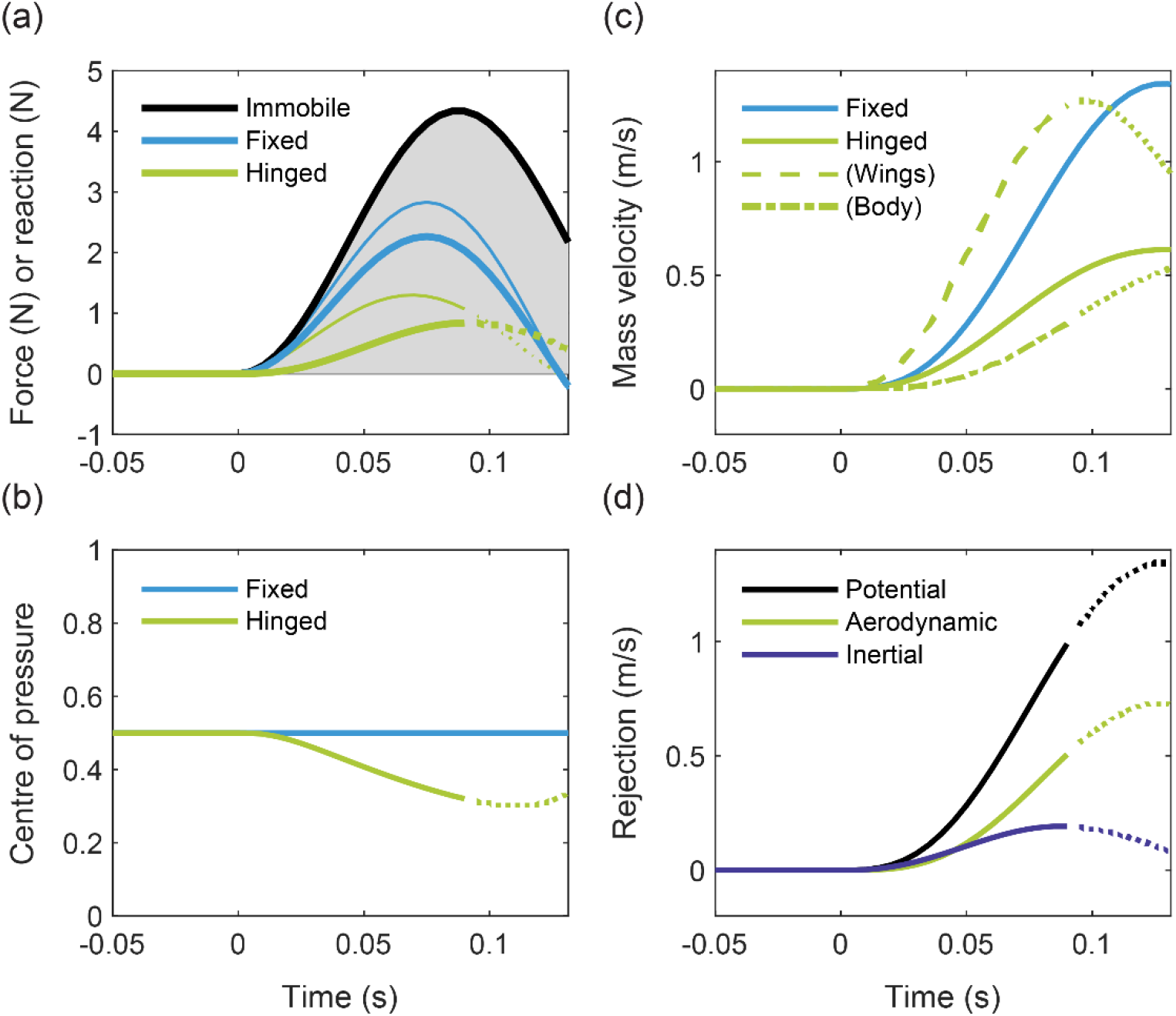
Gust response of the hinged, fixed and immobile systems with an LLC in the 30% gust. Plot lines become dotted if/when the wing angle crosses 20 degrees (approximate onset of nonlinearity). (a) External force Δ*F* = *F* < *F*_0_ (thinner lines) on each wing versus dynamic fuselage reaction Δ*R* (thicker lines). In each case, dynamic reaction is 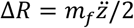 (hinged), Δ*R* = *μ_f_*Δ*F* (fixed) and Δ*R* = Δ*F* (immobile). The gust velocity profile (shaded grey) is shown for reference. (b) The normalised centre of pressure (*l_F_/l*) on each wing. That on the hinged wing moves inboard at first, departing from *P.* (c) Vertical system velocity 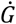 of the hinged (plus component masses, broken lines) and fixed-wing systems. The ascending wings control fuselage motion. (d) Rejection terms for the hinged system. The potential line indicates the maximum achievable rejection, *i.e.,* the amount necessary to keep the fuselage perfectly level.

The transmission of reaction force is successfully delayed in the hinged case. As the centre of pressure drifts inboard under the action of the relative down-flow gradient (Figure 3b), though, alignment with the centre of percussion is progressively lost and fuselage reaction soon develops. The fixed wing, by contrast, transmits load instantly because its fuselage reaction simply scales with the external force (Δ*R* = *μ_f_*Δ*F* applies).

Cheney *et al.* (2020) coined the term *inertial rejection* for the idea that moving wing mass can act to stabilise the fuselage. It offers a complimentary perspective for occasions when forces are unknown or immeasurable—in a kinematics experiment like theirs, for example. Formally, they defined inertial rejection as the vertical velocity difference between the system centre of mass and that of the fuselage, *i.e.,* as the degree of relative fuselage motion or ‘activity’ produced by the wings as they hinge (see SI section 8). Inertial rejection functions best when the fuselage is made to shift by *just* the amount necessary to offset the perturbed system centre of mass, and this only happens when the gust force acts squarely through the centre of percussion. As such, ‘perfect’ inertial rejection *is* the percussion effect. Cheney *et al.* (2020) also introduced the term *aerodynamic rejection* for the difference between the external force upon (or motion of) the hinged and fixed-wing systems. In other words, it is a comparison of the aerodynamic control between the two (note that Cheney *et al.* used vertical velocity from their kinematics data, equivalent to impulse). Altogether, the idea is that inertial rejection lessens the *internal* reaction on the fuselage, while aerodynamic rejection modulates the *external* force on the whole system (via wing morphing or other mechanisms, including relative down-flow). The overall motion is thereby controlled. Indeed, without a reduction in the external force to complement and/or follow inertial rejection, the system would need to rely solely on natural damping from relative down-flow to arrest any acquired motion.

Figure 3c shows how the wings accelerate upwards to absorb the initial brunt of the gust. Inertial rejection is positive but by no means optimal (Figure 3d). Aerodynamic rejection is comparable in magnitude at first, only increasing once the effect of relative down-flow on the hinging wing exceeds that on the fixed, as described. This does not match qualitatively the owl data in Cheney *et al.* (2020), for which inertial rejection is dominant initially.

### 3.3 Rejection by soft stall

Broadly speaking, low-speed aerofoils stall in one of three ways (McCullough & Gault 1951). Two of these precipitate from the leading edge, and the other from the trailing edge. The latter trailing-edge type is associated with thicker sections and is often characterised by a lift curve whose linear region ends smoothly without a sudden drop in lift. Such a curve is sometimes said to exhibit ‘soft stall’ (Eppler 1978; 1990). This contrasts with ‘hard stall’, where lift falls abruptly and often unfavourably. Some soft-stall aerofoils are even designed to have lift curves that reach a maximum and stay there (Wortmann 1972). Their curves plateau, or move onto a region of shallow decline, where lift is largely insensitive to the angle of attack. It transpires that lift curves of this type could enhance the percussion effect by passive means.

We devised a simple approximation for a soft-stall curve, called the *nonlinear lift curve* (NLC), that has a perfect stall plateau (Figure 4a). The plateau begins at *α* = 1/(2*π*)^*c*^ ≈ 9 degrees and *c_L_* = 1 (these values derive from the low-speed sections in Selig *et al.* (1995) at bird-scale Reynolds numbers). We tested the system response with the NLC in the same 30% gust (Figure 4). The gust increases the angle of attack as before, but now the lift coefficient is pushed quickly onto the stall plateau. This happens first at the root, where the linear wing speed is low, and then progressively towards the retreating tip. The force distribution soon approaches saturation (Figure 4b). Not only does this limit the resultant force on each wing, but also it keeps the centre of pressure near the centre of percussion for longer, reducing the initial transmission of reaction to the fuselage (Figure 4c-e). We also plot the linear lift curve (LLC) and fixed-wing NLC data for comparison, which clarify the moment at which lift saturation begins. Notice that the hinged wing with an LLC performs no better than a fixed wing with an NLC; it is the *combination* of the hinge and NLC that works best.

**Figure 4.**
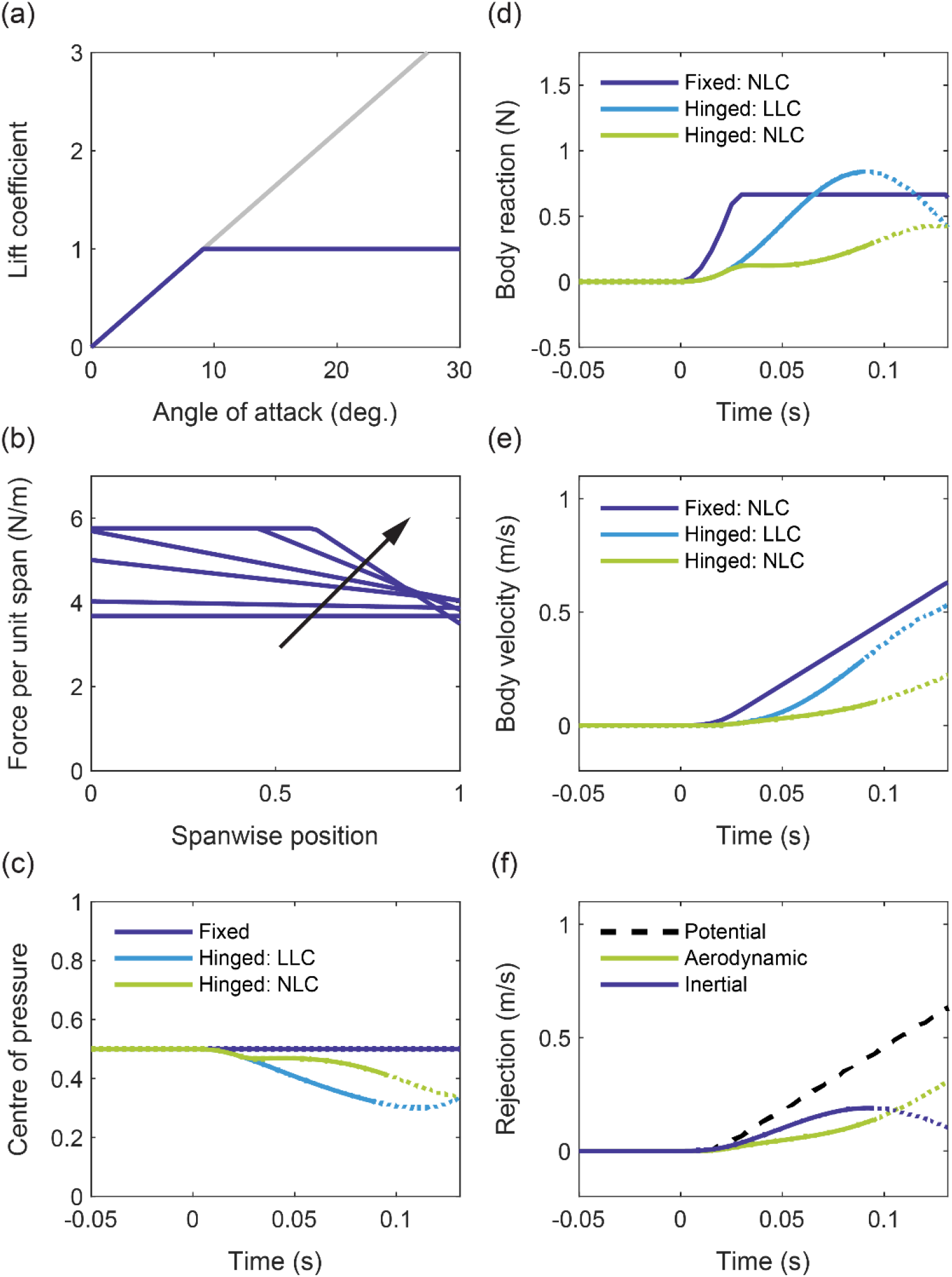
Gust response of the hinged and fixed systems with an L/NLC in the 30% gust. Plot lines become dotted if/when the wing angle crosses 20 degrees (approximate onset of nonlinearity). (a) The NLC lift coefficient is capped at unity. (b) Spanwise force distributions at six evenly spaced instants if time between gust onset (*t* = 0 ms) and peak velocity (*t* = *L_g_*2*U* = 0.0875 ms) for the NLC hinged system. The arrow identifies forward chronology as the force distribution flattens out. Note that the spanwise position has been normalised. (c) Normalised centre of pressure *l_F_/l*. (d) Dynamic fuselage reaction Δ*R.* The departure from the LLC line coincides with the onset of lift saturation. (e) Resulting fuselage velocity 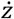. (f) Rejection terms for the NLC hinged system. The potential line indicates the maximum achievable rejection (that necessary to keep the fuselage perfectly level).

Inertial rejection now precedes the aerodynamic (Figure 4f). Both curves bear a compelling resemblance to corresponding data in Cheney *et al.* (2020) for the owl, whose CFD-derived lift curve (for the whole bird) also exhibits soft-stall behaviour. In fact, many birds’ wings also stall gradually at typical flight Reynolds numbers (Withers 1981; van Oorschot 2016 *et al.*) without an abrupt loss of lift. We cannot say for certain whether the resemblance between our data and those in Cheney *et al.* is explained entirely by the lift curve, but soft stall does elicit a strong, sustained percussion effect in our model, and even provides an explanation for delayed aerodynamic rejection (the stalled lift on hinged and fixed wings is similar at first). In any case, the lift curve is undoubtedly important to the dynamics of hinging wings.

### 3.4 Increasing the wing mass fraction

Each hinged wing has thus far made up 10% of the total system mass (Table 1). By increasing the wing mass fraction *μ_w_*, we can enhance the rejection benefit from soft stall. Figure 5 shows the system dynamics for *μ_w_* = 0.35 and *μ_w_* = 0.5, alongside the existing case (*μ_w_* = 0.2). As the relatively heavy wings present greater resistance to motion, less relative down-flow develops across them when they are gusted. Soft stall therefore happens more readily, and the centre of pressure is stabilised accordingly (Figure 5a). Indeed, for the wing of highest mass, the lift distribution reaches total saturation: its resultant effectively behaves as unmoving point force, capped in magnitude and fixed at the halfway mark (*l*/2). The fuselage velocity does not change once this begins (Figure 5b).

**Figure 5.**
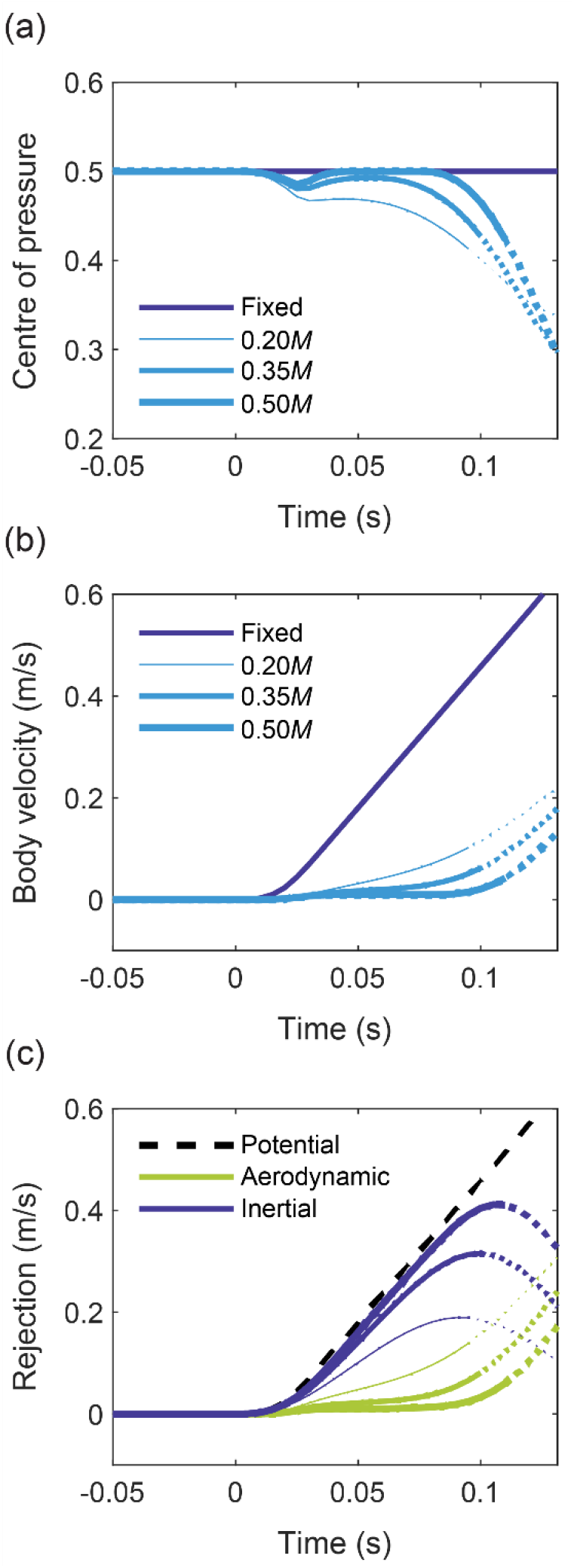
Gust response of the hinged system for three different wing mass fractions, alongside the fixed system, with an NLC in the 30% gust. Line thickness denotes the mass of the hinged wing, from 0.2*M* to 0.5*M.* Plot lines become dotted if/when the wing angle crosses 20 degrees (approximate onset of nonlinearity). (a) Normalised centre of pressure *l_F_/l.* Notice that, with increased mass, the wing is slower to exceed the linear threshold. The model therefore captures the initial dynamics even better. (b) Resulting fuselage velocity 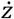. (c) Rejection terms for the hinged system. The higher the relative wing mass, the greater the inertial rejection at first.

Initial aerodynamic rejection falls with increasing *μ_w_* because the fixed and hinged wings each stall quickly with similar total force (Figure 5c), *i.e.*, their CoM velocities are comparable. Inertial rejection, on the other hand, increases with *μ_w_* and becomes the dominant component.

### 3.5 Hinge torque

Hinge torque governs how well inertial rejection works. Unless the torque is well tuned, the wing will not move correctly, no matter where the centre of percussion lies.

The minimum allowable torque for steady gliding flight is *T*_*h*0_, without which the wings would fold up. At low wing angles, a hinge that *exactly* maintains *T*_*h*0_ allows the wing to behave ‘freely’ because static loads cancel, *i.e.*, their moments balance and offer no resistance (or assistance) to motion as the wing rotates. The percussion effect is purest in this constant-torque state, and equation (5) applies. However, a real gliding aircraft cannot fly with constant hinge torque at all times. It must adjust this torque, probably asymmetrically, to the varied demands of flight, including basic gust recovery (restoring the wings to a neutral position) and more advanced control.

In this model, torque can be modulated via parameter Δ*T_h_,* which adds or subtracts from the steady value *T*_*h*0_. If retained in the derivation for the reaction formula, equation (5), dynamic torque gives rise to its own term, or (SI section 6)

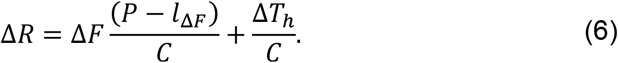

The effect on the fuselage is complicated because the summed terms can interact with each other, with dynamic torque affecting wing motion (hence aerodynamic load) and *vice versa.* To describe the net effect, we define a dynamic centre of percussion 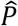 by setting Δ*R* = 0 and solving for *l*_Δ*F*_. Thus

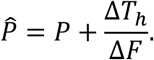

Point 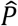 translates along the wing in accordance with the applied force and hinge torque, and if Δ*T_h_* = 0, then 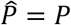 regardless. It is now possible to recast equation (6) as

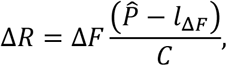

which has the same form as equation (5). The condition of zero reaction now requires that the centre of pressure be kept in alignment with the *dynamic* centre of percussion.

Consider the introduction of a linear spring at the hinge, which adds some static stability to the wing mass. This is perhaps the simplest form of dynamic torque and illustrates well the sensitivity of inertial rejection to the properties of the hinge. Figure 6 shows the effect of various linear torsional springs (*k_t_* = 0.1, 1, 10 N m/rad) on the NLC system, alongside the existing case (*k_t_* = 0 N m/rad) for *μ_w_* = 0.5. As the wing enters the gust front, and the spring engages under extra positive (upward) aerodynamic load, quotient Δ*T_h_*/Δ*F* increases and 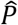 travels outboard from its starting position at *P.* Zero reaction demands that the force move with it; but the NLC tends to prevent this by maintaining, or moving inboard, the centre of pressure on ascending wings (also true for the LLC). Reaction therefore develops as 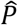 diverges from the centre of pressure, pushing the fuselage upwards. The stiffer the spring, the quicker this happens (Figure 6a), and in the limit of infinite stiffness the system responds exactly as it would with fixed wings. Springs therefore limit inertial rejection by impeding wing motion. The same is true for other mechanical elements that add stiffness or develop resistive torque, including dampers.

**Figure 6.**
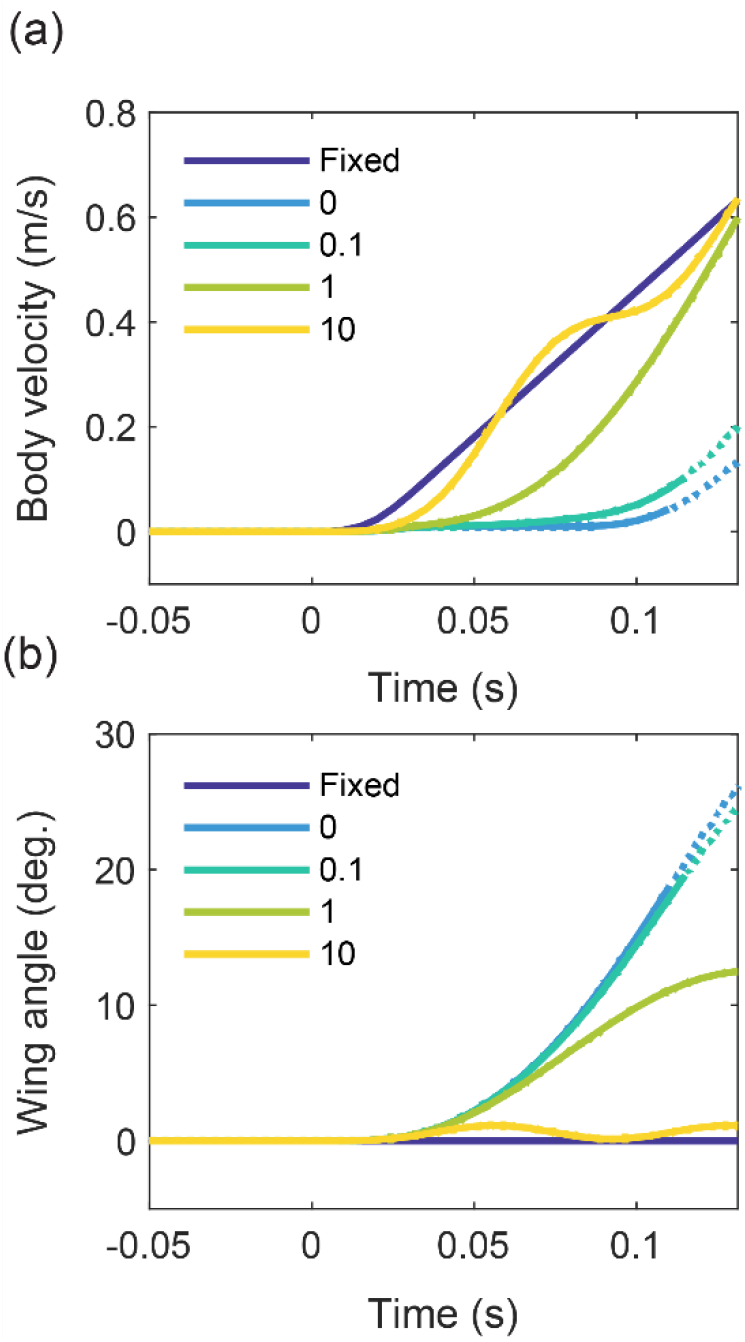
Gust response of the hinged system for three different hinge stiffnesses, versus the fixed system, with an NLC in the 30% gust. Plot lines become dotted if/when the wing angle crosses 20 degrees (approximate onset of nonlinearity). Dynamic torque comes from a linear torsional spring, for which Δ*T_h_* = *k_t_θ*. The legend gives the stiffness constant *k_t_* (N m/rad) for each case. (a) Fuselage velocity 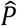. (b) Wing angle *θ.* For the stiffest hinged system, oscillations begin as the lift coefficient hits the stall plateau. This case is illustrative; no well-tuned inertial rejection system would be so stiff or permitted to oscillate in this way.

## 4 Discussion

We show how hinging wings can absorb the initial impact of wing perturbations via the basic mechanics of inertial rejection and the percussion effect. We then propose a method to reject upgusts, which requires: (i) preliminary alignment of the centres of pressure and percussion at equilibrium, *i.e.,* appropriate lift and mass distributions; (ii) hinges that produce constant initial torque; and (iii) wings whose sections stall readily, but softly, onto a lift plateau. Relatively heavy wings are also preferable. Finally, we find that linear spring stiffness at the hinge restricts the motion of the gusted wing, and with it the rejection benefit. Systems with ever stiffer hinges eventually behave as though they had fixed wings (see also Webb & Costello 2008; Oduyela & Slegers 2014).

### 4.1 Mechanics of the hinged wing

Successful gust rejection necessitates a hinge mechanism that can supply the necessary static *and* dynamic torques. The specific requirements are these: (i) support of the static flight loads; (ii) acceptance of the initial motion of the gusted wing with correctly tuned torque, such that the fuselage is isolated from the disturbance; (iii) gradual arrest of the wing while gust load is modulated via aerodynamic rejection; and (iv) prompt restoration of the neutral wing configuration without jolting the fuselage. We leave the detailed implementation of such a hinge for future work, but acknowledge here that the problem is broadly analogous to the design of a suspension system for a terrestrial vehicle. The wings are equivalent to the wheels (the unsprung masses), and the fuselage to the cabin (the sprung mass).

An ideal suspension system provides both *ride quality* and *holding ability.* Ride quality results from isolation of the sprung mass (fuselage or cabin) from the disturbance, and requires appreciable deflection of the unsprung masses (wings or wheels). Holding ability concerns handling and stability, and functions better when deflection of the unsprung masses is limited. As such, these are conflicting criteria (Fischer & Isermann 2004). Basic passive spring-damper suspension systems cannot resolve this conflict and must usually compromise on both counts (this may exclude them from consideration in the proposed rejection method, for which the hinge must work ‘freely’ at first, then in decisive nonlinear fashion). An adaptive active-passive hybrid or fully active suspension system, on the other hand, could provide a better overall solution. These have been commonplace in automotive applications for decades (Sharp & Crolla 1987). Consider, for example, a wing hinge with an active mode that can modify its torque to the instantaneous demands of inertial gust rejection. The hinge would behave passively at first, buying time for the active mode, or, in a *fully* active setup, be driven by a particularly powerful actuator that can mimic the necessary ‘passive’ dynamics. Other measures, such as control-surface deflection, could then provide follow-up aerodynamic rejection. Active methods also permit real-time tuning of the hinge for other flight objectives (Ol *et al.* 2008), including atmospheric energy harvesting, and even allow for adjustments to the dihedral angle(s) for adaptive lateral stability. Ultimately, the designer must decide whether the versatility of hybrid/fully active systems justifies the inevitable mechatronic complexity, extra weight and power demand.

For the system with constant-torque hinges, rejection begins with alignment of the centres of pressure and percussion. This requires suitable distributions of wing lift and mass at equilibrium. On most conventional finite wings, however, the spanwise centre of pressure will naturally lie somewhere near or inside the halfway mark—elliptical loading gives 0.42*l*, for example—and may be difficult to modify without radical alterations to the basic planform or twist geometry. Designers should expect to tune the mass distribution of the wing instead, weighting it towards the hinge for a favourable centre of percussion. The triangular/linear mass distribution is a good starting point; it is realistic, given the usual requirement for structural thickness near the wing root, and might even be achieved by clever placement of electronics, fuel, batteries, or other onboard items.

### 4.2 Stall aerodynamics

Stall is governed by the evolution of the boundary-layer with angle of attack, which itself depends chiefly on the Reynolds number and the shape of the section(s) that make up the wing (McCullough & Gault 1951). Soft stall, in particular, can be achieved by sculpting a wing section to have a surface pressure distribution that slows down the movement of the boundary-layer transition region with incidence (Wortmann 1972; Eppler 1978) for favourable separation behaviour. Many extant wing sections stall softly: Selig *et al.* (1995) provide several conventional designs that do so at bird-scale Reynolds numbers (< 10^5^), while data from Schmitz (1967) show similar behaviour for the simple flat plate. However, the extent to which any of these maintains its soft-stall behaviour during the gust will depend on the timing of the event (the *reduced frequency*) and the associated boundary-layer dynamics. Fast, extreme changes in the flow, including steep gusts, may cause a *dynamic stall* that pushes boundary-layer separation to an angle of attack some way beyond the usual steady value, thereby extending the linear region of the lift curve (McCroskey 1981). The curve will then resemble the LLC of section 3.2. This is an important, open question that motivates further research on low-speed wing sections for optimal stall, including the potential role of boundary-layer control, *e.g.,* suction, blowing or surface-mounted devices (Hazen 1968).

Wings of relatively high mass (section 3.4) have greater rotary inertia and pivot less easily when gusted. The relative down-flow from acquired motion, which acts to oppose the effect of the upgust, is therefore weaker, and soft stall has a better opportunity to develop. If this happens everywhere across the wing, soft stall stabilises the position of the lift vector and extends the percussion effect, buying even more time for other corrective actions to initialise. The benefit is appreciable; near-zero fuselage reaction lasts for ~100 ms in the case of the heaviest wing, long enough for a control system to sense and react to the disturbance. Of course, operating at a high equilibrium angle of attack *α*_0_ (6 degrees here) also facilitates stall.

### 4.3 Avian gust rejection

Birds’ wings, despite their diversity in planform (Lilienthal 1889; Baliga *et al.* 2019) and structural complexity, have spanwise mass distributions that are broadly triangular, becoming higher nearer the shoulder, with local peaks at the elbow and wrist (van den Berg & Rayner 1995; Hedrick *et al.* 2004; Durston 2019). As such, the anatomy naturally bears a mass distribution that puts the centre of percussion near the halfway mark. Now, assuming the lift distribution on these wings is broadly elliptical (exact elliptical lift puts the centre of pressure at 0.42*l*), then close equilibrium alignment between the centres of pressure and percussion may be quite widespread among species—particularly the gliding birds who stand to benefit most from inertial rejection. Of course, the shoulder must be sufficiently compliant, whatever the mass distribution, otherwise the mechanics cannot work at all.

## 5 Conclusions

We present an aeromechanics model of the response of a bird-scale gliding aircraft to a strong, wide upgust. Unlike conventional aircraft, this one has wings that are fully hinged to the fuselage on pin joints that enable rotation in the vertical plane. The hinged design was inspired by the response of birds to upgusts, as measured in a laboratory experiment.

Hinging allows the perturbed wings to absorb and reject the brunt of a disturbance. The rejection can be optimised by having two key spanwise points on the wing, the centres of *pressure* and the *percussion*, start and stay in good alignment during these early moments of the gust. Initial transmission of load to the fuselage is thereby delayed and/or reduced (which would buy time for other flight control processes to initialise). We call this the ‘percussion effect’. Having presented the basic mechanics, we propose a passive method for achieving the effect in upgusts. The essential ingredients are: (i) well-considered lift and mass distributions for equilibrium alignment of the two key points; (ii) hinges with constant initial torque output (enough for aircraft weight support but no more or less); and (iii) a wing whose sections stalls softly, such that the centre of pressure is stabilised during gusted rotation.

We ultimately envision the mechanics of the percussion effect as part of a complete hinged-wing suspension system, primarily for small aircraft operating in the gusty conditions of the low atmosphere.

## Acknowledgements

The authors wish to thank Ronald C. M. Cheung and Amir K. Bagheri for insightful discussions on the modelling.

## References

Baliga, V. B., Szabo, I. and Altshuler, D. L. (2019) ‘Range of motion in the avian wing is strongly associated with flight behavior and body mass’, Science Advances, 5 (10).

Brearley, M. N., Burns, J. C. & De Mestre, N. J. (1998) ‘What is the best way to hit a cricket ball?’, International Journal of Mathematical Education in Science and Technology, 21 (6), pp. 949–961.

Cheney, J. A., Stevenson, J. P. J., Durston, N. E., Song, J., Usherwood, J. R., Bomphrey, R. J. and Windsor, S. P. (2020) ‘Bird wings act as a suspension system that rejects gusts’, Proceedings of the Royal Society B, 287 (1937).

Cross, R. (1998) ‘The sweet spot of a baseball bat’, American Journal of Physics, 66 (9), pp. 772–779.

Durston, N. (2019) Quantifying the flight stability of free-gliding birds of prey, PhD Thesis, University of Bristol.

Eppler, R. (1978) ‘Turbulent Airfoils for General Aviation’, Journal of Aircraft, 15 (2).

Eppler, R. (1990) Airfoil Design and Data, Berlin: Springer-Verlag.

Etkin, B. (1981) ‘Turbulent Wind and Its Effect on Flight’, Journal of Aircraft, 18 (5).

Federal Aviation Administration (FAA) (2020) ‘Gust and turbulence loads’, in Code of Regulations (Title 14, Chapter 1, Subchapter C, Part 25, section 341), Washington D. C.: Office of the Federal Register.

Fischer, D. and Isermann, R. (2004) ‘Mechatronic semi-active and active vehicle suspensions’, Control Engineering Practice, 12 (11), pp. 1353–1367.

Fung, Y. C. (1993) An Introduction to the Theory of Aeroelasticity, New York: Dover.

Gray, J. (1968) Animal Locomotion, London: Weidenfeld & Nicolson.

Hazen, D. C. (1968) ‘Film Notes for Boundary-layer Control’, National Committee for Fluid Mechanics Films: Cambridge.

Hedrick, T. L., Usherwood, J. R. and Biewener, A. A. (2004) ‘Wing inertia and whole-body acceleration: an analysis of instantaneous aerodynamic force production in cockatiels (*Nymphicus hollandicus*) flying across a range of speeds’, Journal of Experimental Biology, 207, pp. 1689–1702.

Leylek, E. A. and Costello, M. (2015) ‘Use of Compliant Hinges to Tailor Flight Dynamics of Unmanned Aircraft’, Journal of Aircraft, 52 (5).

Lilienthal, O. (1889) Der Vogelflug als Grundlage der Fliegekunst, Berlin: Hermann Heyfelder.

Lissaman, P. B. S. (1983) ‘Low-Reynolds-Number Airfoils’, Annual Review of Fluid Mechanics, 15, pp. 223–239.

McCroskey, W. J. (1981) The Phenomenon of Dynamic Stall, NASA-TM-81264, Moffett Field: NASA.

McCullough, G. B. & Gault, D. E. (1951) Examples of Three Representative Types of Airfoil-section Stall at Low Speed, NACA TN 2505, Moffett Field: NACA.

Oduyela, A. and Slegers, N. (2014) ‘Gust Mitigation of Micro Air Vehicles Using Passive Articulated Wings’, The Scientific World Journal, 2014.

Ol, M., Parker, G., Abate, G. and Evers, J. (2008) ‘Flight Controls and Performance Challenges for MAVs in Complex Environments’, AIAA Guidance, Navigation and Control Conference and Exhibit, Honolulu, 18-21 August, Reston: AIAA.

Paranjape, A. A., Chung, S-J. and Hilton, H. H. (2012) ‘Dynamics and Performance of Tailless Micro Aerial Vehicle with Flexible Articulated Wings’, AIAA Journal, 50 (5).

Rao, S. S. (2004) Mechanical Vibrations, 4^th^ edn, Pearson Prentice Hall: Hoboken.

Reynolds, K. V., Thomas, A. L. R. and Taylor, G. K. (2014) ‘Wing tucks are a response to atmospheric turbulence in the soaring flight of the steppe eagle *Aquila nipalensis*’, Journal of the Royal Society Interface, 11 (101).

Sabins, R. (1937) Flexible Airplane Wing Construction, US Patent Office, Patent no. 2066649.

Schmitz, F. W. (1967) Aerodynamics of the Model Airplane. Part 1. Airfoil Measurements, Translated by Redstone Scientific Information Center (RSIC), Redstone Arsenal: RSIC.

Selig, M. S., Guglielmo, J. J., Broeren, A. P. and Giguère, P. (1995) Summary of Low-Speed Airfoil Data, Virginia Beach: SoarTech Publications.

Sharp, R. S. and Crolla, D. A. (1987) ‘Road Vehicle Suspension System Design – a review’, Vehicle System Dynamics, 16 (3), pp. 167–192.

Stewart, K. C., Blackburn, K., Wagener, J., Czabaranek, Lt. J. and Abate, G. (2008) ‘Development and Initial Flight Tests of a Single-Jointed Articulated-Wing Micro Air Vehicle’, AIAA Atmospheric Flight Mechanics Conference and Exhibit, Honolulu, 18-21 August, Reston: AIAA.

Tennekes, H. (2009) The Simple Science of Flight, Cambridge: MIT press.

van den Berg & Rayner, C. and Rayner, J. (1995) ‘The moment of inertia of bird wings and the inertial power requirement for flapping flight’, Journal of Experimental Biology, 198, pp. 1655–1664.

van Oorschot, B. K., Mistick, E. A. and Tobalske, B. W. (2016) ‘Aerodynamic consequences of wing morphing during emulated take-off and gliding in birds’, Journal of Experimental Biology, 219, pp. 3146–3154.

Watkins, S. W., Milbank, J., Loxton, B. J. and Melbourne, W. H. (2006) ‘Atmospheric Winds and Their Implications for Microair Vehicles’, AIAA Journal, 44 (11).

Webb, A. and Costello, M. (2008) Wing Articulation of Micro Air Vehicles to Reduce Gust Sensitivity, Atlanta: Georgia Institute of Technology.

Wilson, T., Kirk, J, Hobday, J. and Castrichini, A. (2019) ‘Small scale flying demonstration of semi aeroelastic hinged wing tips’ (IFASD-2019-076), International Forum on Aeroelasticity and Structural Dynamics, Savannah, 9-16 June.

Withers, P. C. (1981) ‘An aerodynamic analysis of bird wings as fixed aerofoils’, Journal of Experimental Biology, 90, pp. 143–162.

Wortmann, F. X. (1972) ‘A Critical Review of the Physical Aspects of Airfoil Design at Low Mach Numbers’, in Motorless Flight Research (NASA CR-2145), Washington D. C.: NASA.

Wright, J. R. and Cooper, J. E. (2007) Introduction to Aircraft Aeroelasticity and Loads, Chichester: Wiley.

