## supplemental_material for "Dynamics of hinged wings in strong upward gusts"

Expressions in dark orange are featured in the main manuscript.

### SI section 1 Equations of motion for the hinged system

We derived the equations of motion for the hinged system by Lagrange's energy method. Lagrange's equations, in compact indicial notation, are

$$\frac{\mathrm{d}}{\mathrm{d}t}\left(\frac{\partial L}{\partial \dot{q}_i}\right)-\frac{\partial T}{\partial q_i}=Q_i,$$

where $L = T - V$ is the *Lagrangian*, the difference between the kinetic ($T$) and potential ($V$) energies of the system, and $Q_i$ are the generalised, non-conservative loads. Index $i$ runs across each generalised coordinate or degree of freedom ($z$ and $\theta$ in this case). By substitution and expansion,

$$\frac{\mathrm{d}}{\mathrm{d}t}\left(\frac{\partial T}{\partial \dot{q}_i}\right)-\frac{\partial T}{\partial q_i}+\frac{\partial V}{\partial q_i}=Q_i.$$

Note that we set

$$\frac{\mathrm{d}}{\mathrm{d}t}\left(\frac{\partial V}{\partial \dot{q}_i}\right)=0$$

because potential energy $V$ is not a function of time or velocity in mechanical systems. It remains to find expression for the energies and loads, then to evaluate the Lagrange equation for each coordinate. The total kinetic energy of the system is

$$T=\frac{M\dot{z}^2}{2}+2m_w\dot{z}l_m\dot{\theta}\cos\theta+I_h\dot{\theta}^2,$$

and the total potential energy is

$$V = Mgz + 2m_w g l_m \sin\theta.$$

The generalised loads are found by considering the work done by each non-conservative force and/or torque along/through each coordinate direction. Along $z$ (fixing $\theta$), the generalised force is

$$Q_z = 2F\cos\theta,$$

and through $\theta$ (fixing $z$), the generalised torque is

$$Q_\theta = 2Fl_F - 2T_h.$$

We can now find each equation of motion. For $z$, the Lagrange equation is

$$\frac{\mathrm{d}}{\mathrm{d}t}\left(\frac{\partial T}{\partial \dot{z}}\right)-\frac{\partial T}{\partial z}+\frac{\partial V}{\partial z}=Q_z,$$

which, after substitution and simplification, gives

$$M\ddot{z}+2m_w l_m\left(\ddot{\theta}\cos\theta-\dot{\theta}^2\sin\theta\right)+Mg=2F\cos\theta.$$

Similarly, the Lagrange equation for $\theta$ is

$$\frac{\mathrm{d}}{\mathrm{d}t}\left(\frac{\partial T}{\partial \dot{\theta}}\right)-\frac{\partial T}{\partial \theta}+\frac{\partial V}{\partial \theta}=Q_\theta,$$

ultimately giving

$$I_h\ddot{\theta}+m_w l_m\ddot{z}\cos\theta+T_h+m_w g l_m\cos\theta=Fl_F.$$

Each equation can also be obtained by direct application of Newton's laws. The first is the 2nd law for vertical motion of the *system* centre of mass, and the second the same law but for torques/moments on one wing.

**SI section 2 Subtraction of static loads**

The linearised equations of motion for the hinged system are (with small-angle assumptions)

$$M\ddot{z} + 2m_w l_m \ddot{\theta} + Mg = 2F$$

and

$$I_h \ddot{\theta} + m_w l_m \ddot{z} + T_h + m_w g l_m = F l_F.$$

We can subtract the static loads from these equations to leave only increments. This also removes the influence of gravity. Before the gust strikes, all time derivatives vanish, and every other term assumes its equilibrium value (subscript 0). We obtain the equations of *static equilibrium*,

$$Mg = 2F_0$$

and

$$T_{h0} + m_w g l_m = F_0 l_{F0},$$

for the condition of steady level flight. Upon subtraction of these static loads,

$$M\ddot{z} + 2m_w l_m \ddot{\theta} = 2(F - F_0)$$

and

$$I_h \ddot{\theta} + m_w l_m \ddot{z} + (T_h - T_{h0}) = (F l_F - F_0 l_{F0}).$$

The bracketed terms now contain the difference between total and equilibrium values. We denote these *dynamic* increments by $\Delta$. For example, $\Delta F$ is the vertical force increment on each wing. Thus,

$$M\ddot{z} + 2m_w l_m \ddot{\theta} = 2\Delta F$$

and

$$I_h \ddot{\theta} + m_w l_m \ddot{z} + \Delta T_h = \Delta F l_{\Delta F}.$$

On the right-hand side: it is perhaps not immediately obvious that $F l_F - F_0 l_{F0}$ should become $\Delta F l_{\Delta F}$. This is just a restatement of the centroid principle, *i.e.*, that the total moment ($F l_F$) is the sum of its constituents, both static ($F_0 l_{F0}$) and dynamic ($\Delta F l_{\Delta F}$).

### SI section 3 Wing mass properties

The triangular wing mass distribution has the form

$$m_w' = Ay + B.$$

Coordinate $y$ begins at the wing root and points outboard along the length. Coefficients $A$ and $B$ are found by applying two boundary conditions, $m_w' = 0$ at $y = l$, and

$$m_w = \int_0^l m_w' \, \mathrm{d}y$$

(total mass). Hence, we find $B = -Al$ and $A = -2m_w/l^2$, so that

$$m_w' = -\frac{2m_w}{l}\left(\frac{y}{l} - 1\right).$$

The inertial properties of this distribution may be found by integration. The centre of mass $l_m$ derives from

$$m_w l_m = \int_0^l m_w' y \, \mathrm{d}y,$$

which gives $l_m = l/3$. The moment of inertia is

$$I_h = \int_0^l m_w' y^2 \, \mathrm{d}y$$

which gives $I_h = m_w l^2/6$. The radius of gyration is then $k_w = \sqrt{I_h/m_w} = l/\sqrt{6}$. The centre of percussion is given by (derived in section 6)

$$P = \frac{I_h}{m_w l_m}\left(= \frac{k_w^2}{l_m}\right),$$

which, in this case, gives $P = l/2$. The centre of percussion can also be expressed as

$$P = \frac{\int_0^l m_w' y^2 \, \mathrm{d}y}{\int_0^l m_w' y \, \mathrm{d}y},$$

*i.e.*, the quotient of the 2nd and 1st moments of the mass distribution about the hinge.

The relative placement of the key inertial points ($l_m$, $k_w$ and $P$) can be deduced by simple reasoning. It must be true that

$$P \geq l_m,$$

otherwise the perturbation force cannot produce the necessary moment, thence rotary motion, about the centre of mass to oppose the translation of the hinge. We now construct two inequalities,

$$l_m = \frac{k_w^2}{P} \leq P$$

and

$$P = \frac{k_w^2}{l_w} \geq l_m.$$

These give $P^2 \geq k_w^2 \geq l_m^2$ or $P \geq k_w \geq l_m$. $P$ is therefore outermost and $l_m$ innermost, with $k_w$ in between.

**SI section 4 Aerodynamics formulation**

The approach is fundamentally similar to *blade-element theory*.

(For example, see Houghton, E. L and Carpenter, P. W. (1993) *Aerodynamics for Engineering Students*, 4th edn, London: Hodder & Stoughton.)

$F$ is the aerodynamic force on each wing in the (vertical) plane of motion. It acts at distance $l_F$ from the hinge. In the equations of motion, we define $F$ as perpendicular to the wing; it is therefore the vertical projection of $F$, or $F\cos\theta$, that determines the vertical motion of the system. This projection becomes approximately equal to $F$ itself when wing angle $\theta$ is linearised (Figure SI-4-1 a). All subsequent references to $F$ refer to this projection.

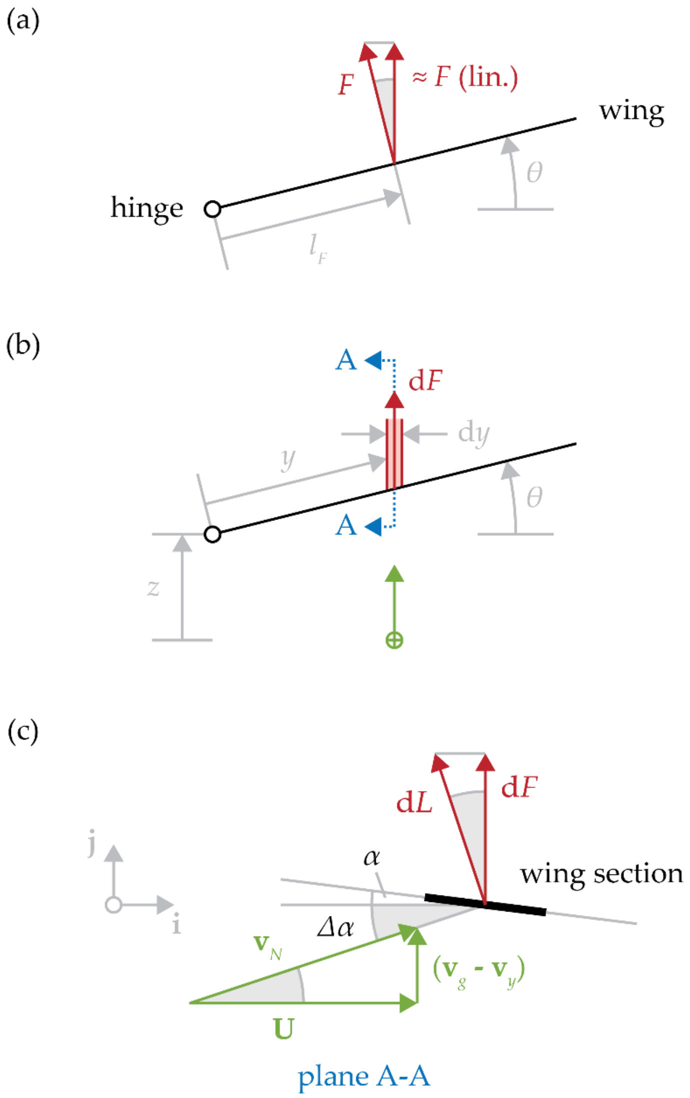


**Figure SI-4-1** Force $F$ on the wing comes from elemental contributions $\mathrm{d}F = F'\mathrm{d}y$. (a) Front view showing $F$ and its linearised (lin.) vertical projection. (b) Front view showing element $\mathrm{d}y$. Gust $\mathbf{v}_g$ spans the entire system, but only that slice of flow acting in plane A-A (green vector) is shown. (c) Side view of the A-A wing section (black line). The shaded angles are equal ($=\Delta\alpha$).

$F$ is the integral resultant from the elemental contributions $\mathrm{d}F = F'\mathrm{d}y$ (also vertical in the linearisation) of the force distribution $F'$ across the wing. In turn, this distributed force depends on the instantaneous flow conditions, including the gust strength and/or induced flow due to motion of the system. We now find expressions for $F$ and its aerodynamic hinge moment $M_F$.

A-A will be the plane of analysis, at spanwise distance $y$ from the hinge. We express the resultant relative wind vector $\mathbf{v}_N$ in the $\mathbf{ij}$ coordinate system, which lies in this plane (Figure SI-4-1 b-c). The vector has three components:

1. Freestream velocity $\mathbf{U}$.
2. Upgust velocity $\mathbf{v}_g = \mathbf{v}_g(t)$.
3. Induced flow velocity $\mathbf{v}_y = \mathbf{v}_y(t)$.

The freestream velocity $\mathbf{U}$ is constant and equal-and-opposite to the forward flight speed. It acts along positive $\mathbf{i}$, so $\mathbf{U} = U\mathbf{i}$.

The (time-varying) upgust velocity $\mathbf{v_g}$ points along positive $\mathbf{j}$, so $\mathbf{v}_g = v_g\mathbf{j}$.

Induced flow arises from motion acquired by the gusted wing. The absolute velocity of the wing section in A-A is

$$\mathbf{v}_A = \mathbf{v}_h + \mathbf{v}_{A/h},$$

*i.e.*, the velocity of the hinge (subscript *h*) plus that of A-A relative to the hinge ($A/h$). The absolute induced flow velocity $\mathbf{v}_y$ is then $-\mathbf{v}_A$. In the linear framework (low $\theta$),

$$\mathbf{v}_y \backsimeq -\left(\dot{z} + y\dot{\theta}\right)\mathbf{j}.$$

Altogether, $\mathbf{v}_N = \mathbf{U} + \mathbf{v}_g + \mathbf{v}_y$, so

$$\mathbf{v}_N = U\mathbf{i} + \left(v_g - \dot{z} - y\dot{\theta}\right)\mathbf{j}.$$

This vector has magnitude

$$v_N^2 = \|\mathbf{v}_N\|^2 = U^2 + \left(v_g - \dot{z} - y\dot{\theta}\right)^2,$$

and modifies the equilibrium section angle of attack $\alpha_0$ by increment $\Delta\alpha$, according to

$$\tan\Delta\alpha = \frac{v_g - \dot{z} - y\dot{\theta}}{U}.$$

Assuming the gust is moderate in strength (see manuscript), we can linearise $\Delta\alpha$ with the approximation $\tan\Delta\alpha \sim \Delta\alpha$. Hence,

$$\Delta\alpha \backsimeq \frac{v_g - \dot{z} - y\dot{\theta}}{U}.$$

As mentioned, force $F$ is the resultant of spanwise distribution $F'$, which is related to the lift distribution $L'$. Consider first the lift distribution (lift per unit wing length),

$$L' = \frac{\rho v_N^2}{2} c c_L(\alpha_0 + \Delta\alpha),$$

where $c$ is the chord length, and $c_L$ the lift coefficient. Since $\mathrm{d}L = L'\mathrm{d}y$, $\mathrm{d}F = \mathrm{d}L\cos\Delta\alpha$ and $v_N^2 = U^2/\cos^2\Delta\alpha$ (Figure SI-4-1 c),

$$\mathrm{d}F = F'\mathrm{d}y = qc\frac{c_L(\alpha_0 + \Delta\alpha)}{\cos\Delta\alpha}\mathrm{d}y,$$

where $q = \rho U^2/2$ is the dynamic pressure. For small changes, $\cos\Delta\alpha \sim \Delta\alpha$. Across the wing length $l$, then,

$$F = qc\int_0^l c_L(\alpha_0 + \Delta\alpha)\,\mathrm{d}y,$$

The corresponding elemental moment from $\mathrm{d}F$ is $\mathrm{d}M_F = y\mathrm{d}F$, so

$$M_F = qc\int_0^l y c_L(\alpha_0 + \Delta\alpha)\,\mathrm{d}y.$$

Each of these effectively contains static (due to $\alpha_0$) and dynamic (due to $\Delta\alpha$) parts. Force $F$, for example, can be broken into $F_0$ and $\Delta F$, where

$$\Delta F = qc \int_0^l c_L(\Delta\alpha)\,\mathrm{d}y.$$

In our calculations, we use discretised versions of these integrals with 50 spanwise points along the length of each wing. Finally, the point of action of force $F$, or its spanwise centre of pressure, is

$$l_F = \frac{M_F}{F}.$$

### SI section 5 Unsteady (build-up) effects

Force on the wings takes time to 'build up' following a change in the flow. We approximated and checked this effect using classical approaches, viz. Küssner and Wagner functions, and compared the results to the quasi-steady response. We used the case of the linear lift curve (LLC) throughout. This section gives the relevant equations; see Fung (1993) and Wright & Cooper (2007) for background details.

First, we transform the equations of motion into dimensionless time $\tau$ ($= 2Ut/c$). Thus,

$$M\left(\frac{2U}{c}\right)^2 z'' + 2m_w l_m \left(\frac{2U}{c}\right)^2 \theta'' = 2\Delta F$$

and

$$I_h\left(\frac{2U}{c}\right)^2 \theta'' + m_w l_m \left(\frac{2U}{c}\right)^2 z'' + \Delta T_h = \Delta M_F.$$

The prime symbol ($'$) indicates differentiation with respect to the new time variable.

For this unsteady model, the lift force ($\Delta F$) and moment ($\Delta M_F$) increments must be modified to include unsteady build-up effects from: (i) the gust; (ii) relative flow from induced motion; and (iii) added (air) mass. First, we consider the lift $\Delta F$.

(i) The modified lift increment (per unit wing length) from the **gust** is

$$\Delta F_g' = aqc \int_0^\tau \frac{v_g(\tau_0)}{U} \Psi'(\tau - \tau_0)\, \mathrm{d}\tau_0.$$

$\Psi$ is the *Küssner function*, and $\Psi'$ its $\tau$-derivative. For infinite (2D) wings at low speed, a good approximation is

$$\Psi = 1 - 0.5e^{-0.13\tau} - 0.5e^{-\tau}.$$

(ii) The modified lift increment (per unit wing length) from **relative flow** in $z$ is

$$\Delta F_z' = 2aq \int_0^\tau h''(\tau_0)\Phi(\tau - \tau_0)\, \mathrm{d}\tau_0,$$

where $h'' = z'' + y\theta''$ is the (linearised) acceleration of the wing section at distance $y$ from the hinge. $\Phi$ is the *Wagner function*, well approximated by

$$\Phi = 1 - 0.165e^{-0.045\tau} - 0.335e^{-0.3\tau}.$$

These are convolution (or *Duhamel*) integrals. Each one convolves an aspect of the changing flow ($v_g/U$ or $h''$) with a sliding modifier function ($\Psi'$ or $\Phi$, in turn). As such, they account for the time history of the flow in the calculation of force.

(iii) The lift increment (per unit wing length) from the **added mass effect** is

$$\Delta F_m' = 2aqh''.$$

It is, however, negligible in this case.

Standard 'star' notation ($*$) will denote convolution. The Küssner integral becomes

$$\Delta F_g' = aqc\frac{(v_g * \Psi')}{U},$$

and the Wagner equivalent,

$$\Delta F_z' = 2aq(h'' * \Phi).$$

The net lift force (per unit wing length) is then $\Delta F' = \Delta F_g' - \Delta F_z'$, neglecting added mass. If we substitute $\Delta F_g'$ and $\Delta F_z'$ and integrate this across the wing length $l$, we obtain the overall wing lift increment,

$$\Delta F = aq\left[\frac{c}{U}\left(v_g * \Psi'\right) - 2\left(z'' + \frac{b}{2}\theta''\right) * \Phi\right].$$

By a similar exercise, we find the wing moment,

$$\Delta M_F = Fl_{\Delta F} = aq\left[\frac{c}{2U}\left(v_g * \Psi'\right) - \left(z'' + \frac{2b}{3}\theta''\right) * \Phi\right].$$

We now transplant these into the equations of motion and apply the Laplace transform, converting the entire system to the $s$ domain for easier solution (in MATLAB). We omit these steps for brevity.

Summary: by switching the build-up functions on and off in turn ('off' means setting a function to unity, *e.g.*, $\Psi = 1$) and comparing the solutions for the state variables, we found that the Küssner effect, pertaining to the gust, is the more important overall. Convolution with the Küssner function lessens (by ~25%) and spreads the input gust profile somewhat, also extending its tail, thereby modifying the perturbation 'seen' by the system. That said, the area under the gust profile—equivalent to the input energy—is conserved post-convolution, and the overall impact on the system dynamics is undramatic. The quasi-steady result is effectively just pushed back in time. We therefore decided to neglect build-up effects in the model proper, favouring instead the simpler, more versatile quasi-steady aerodynamics formulation.

### SI section 6 Linearised reaction formulae

Recall the linearised equations of motion for the hinged system,

$$M\ddot{z} + 2m_w l_m \ddot{\theta} = 2\Delta F$$

and

$$I_h \ddot{\theta} + m_w l_m \ddot{z} + \Delta T_h = \Delta F l_{\Delta F}.$$

By finding an expression for $\ddot{\theta}$ from the first equation and putting it into the second, we can solve for the body acceleration $\ddot{z}$. The first step, once like terms have been collected, gives

$$\ddot{z}\left(m_w l_m - \frac{I_h M}{2m_w l_m}\right) + \Delta T_h = \Delta F\left(l_{\Delta F} - \frac{I_h}{m_w l_m}\right),$$

and we find

$$\ddot{z} = \frac{\Delta F}{C'}\left(l_{\Delta F} - \frac{I_h}{m_w l_m}\right) - \frac{\Delta T_h}{C'},$$

where

$$C' = \left(m_w l_m - \frac{MP}{2}\right).$$

The equation of motion for the body is simply $\Delta R = m_f \ddot{z}/2$. We now solve for reaction $\Delta R$ as

$$\Delta R = \Delta F \frac{(P - l_{\Delta F})}{C} + \frac{\Delta T_h}{C},$$

where

$$P = \frac{I_h}{m_w l_m}$$

is the distance along the wing from the hinge to the *centre of percussion*. Constant $C = P - (\mu_w/\mu_f)(l_m - P)$ is positive and includes inertial/mass properties. Notice that $P$ and $l_{\Delta F}$ have been switched in their brackets, and the signs flipped accordingly.

For the case when dynamic torque is zero, $\Delta T_h = 0$, we find

$$\Delta R = \Delta F \frac{(P - l_{\Delta F})}{C}.$$

For the fixed-wing system, the equation of motion is

$$M\ddot{z} + Mg = 2F.$$

At equilibrium, $Mg = 2F_0$, so $M\ddot{z} = 2\Delta F$. The body follows a similar equation,

$$m_f g + m_f \ddot{z} = 2R,$$

which becomes $m_f \ddot{z} = 2\Delta R$ when static loads are removed. The accelerations the body and system must be the same, so we find $\Delta R = \mu_f \Delta F$.

### SI section 7 Load modulation interval

For the constant-torque hinge, the transmitted reaction is

$$\Delta R = \Delta F \frac{(P - l_{\Delta F})}{C},$$

and for the fixed-wing system, $\Delta R = \mu_f \Delta F$. The question of when the hinged system outperforms the fixed one is therefore answered by solving the inequality

$$\left|\frac{(P - l_{\Delta F})}{C}\right| < \mu_f.$$

(We have cancelled $\Delta F$, which must be the same in a like-for-like comparison.) This breaks into

$$l_{\Delta F} > P - \mu_f C$$

and

$$l_{\Delta F} < P + \mu_f C.$$

Since $C = P - (\mu_w/\mu_f)(l_m - P)$, we can now solve for the interval. Thus,

$$P - (P - \mu_w l_m) < l_{\Delta F} < P + (P - \mu_w l_m),$$

or that $l_{\Delta F}$ must lie within $P \pm (P - \mu_w l_m)$. This interval depends on the mass distribution of the wing.

Note: if $\Delta F$ acts *exactly* on the inner bound, the wing will stay level as it is pushed, as in the fixed case.

**SI section 8 Inertial rejection**

We give here the mathematical link between inertial rejection and wing motion.

Inertial rejection $v_I$ is the difference between the vertical velocity of the system centre of mass and the body, or $\dot{G} - \dot{z}$. First, from the method of mass-moments,

$$MG = zm_f + 2m_w(z + l_m \sin\theta).$$

Since $M = m_f + 2m_w$ and $\mu_w = 2m_w/M$, we find

$$G = z + \mu_w l_m \sin\theta,$$

so

$$\dot{G} = \dot{z} + \mu_w l_m \dot{\theta} \cos\theta.$$

Hence, $v_I = \dot{G} - \dot{z} = \mu_w l_m \dot{\theta} \cos\theta$. Inertial rejection is therefore proportional to the vertical (on account of the cosine) linear velocity of the centre of wing mass relative to the body.
